# ClpAP proteolysis does not require rotation of the ClpA unfoldase relative to ClpP

**DOI:** 10.1101/2020.07.26.221812

**Authors:** Sora Kim, Kristin L Zuromski, Tristan A Bell, Robert T Sauer, Tania A Baker

## Abstract

AAA+ proteases, which perform regulated protein degradation in all kingdoms of life, consist of a hexameric AAA+ unfoldase/translocase in complex with a self-compartmentalized peptidase. Based on asymmetric features of cryo-EM structures and a sequential hand-over-hand model of substrate translocation, recent publications have proposed that the AAA+ unfoldases ClpA and ClpX must rotate with respect to their partner peptidase ClpP to allow function. Here, we test this model by covalently crosslinking ClpA to ClpP to prevent rotation. We find that crosslinked ClpAP complexes unfold, translocate, and degrade protein substrates, albeit modestly slower than uncrosslinked enzyme controls. Rotation of ClpA with respect to ClpP therefore is ot required for ClpAP protease activity, although some flexibility in how the AAA+ ring ocks on ClpP may be necessary for optimal function.

## Introduction

The AAA+ (<u>A</u>TPases <u>A</u>ssociated with diverse cellular <u>A</u>ctivities) protease subfamily uses the energy of ATP binding and hydrolysis to disassemble and degrade proteins that are misfolded, deleterious, or unneeded (Sauer and Baker, 2011). AAA+ proteases are composed of a hexameric single-or double-ringed AAA+ unfoldase/translocase and a self-compartmentalized partner peptidase, which assemble as stacked rings. Upon substrate recognition by the unfoldase, repeated cycles of ATP hydrolysis power conformational changes in the AAA+ motor, promoting substrate unfolding and processive translocation of the resulting polypeptide into the protease chamber for degradation. Recent structural and biochemical studies have illuminated some aspects of this process, but the stepwise mechanisms these proteolytic machines use to carry out mechanical unfolding and translocation of protein substrates are still being explored (Puchades *et al*., 2020).

ClpAP protease is constructed from the double-ring hexameric ClpA6 AAA+ unfoldase and the tetradecameric ClpP14 peptidase, which consists of two heptameric rings (Sauer and Baker, 2011; Figure 1A). Thus, the interface between ClpA and ClpP has an asymmetric six-to-seven subunit mismatch. The ClpXP protease, which consists of the single-ring AAA+ ClpX unfoldase and the ClpP peptidase, has the same six-seven mismatch, as do AAA+ proteasomal enzymes. How such mismatches are accommodated structurally and whether they relate to the mechanisms of these ATP-dependent proteases has long been a subject of interest. Recent cryo-EM structures of ClpAP and ClpXP reveal six flexible loops protruding from the bottom face of the ClpP-proximal AAA+ ring. Each loop contains a conserved tripeptide motif (IGL in ClpA; IGF in ClpX) that docks into five or six of the seven hydrophobic binding clefts on the “top” of the ClpP ring (Fei *et al*., 2020; Ripstein *et al*., 2020; Lopez *et al*., 2020). Intriguingly, the unoccupied ClpP clefts are always located between two specifically positioned subunits in the ClpA or ClpX hexamers (Figure 1A).

**Figure 1.**
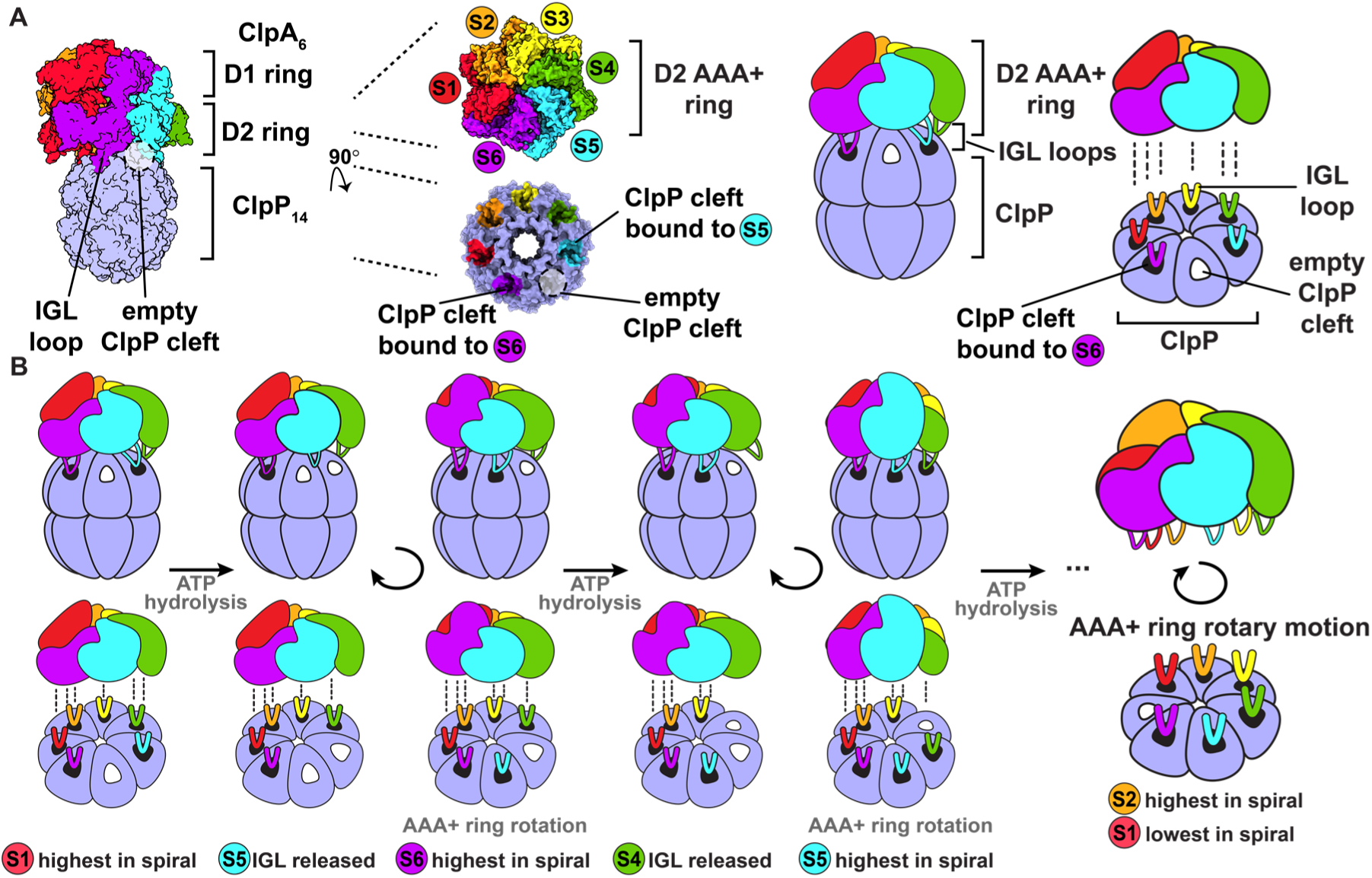
ClpAP structure and rotary translocation model. (**A**) Complex of ClpP with ClpA (PDB 6UQO). Subunits of ClpA, labeled S1 through S6, are ordered from the highest to the lowest position in the spiral relative to ClpP at the beginning of the mechanical cycle. The IGL loops of ClpA hexamers dock into a subset of the seven clefts in a heptameric ClpP ring. There is an empty cleft between the second lowest and lowest subunits in the spiral (S5 and S6, respectively). The coloring of the ClpP clefts represents the docked position of the IGL loops from the corresponding AAA+ subunits; empty clefts are represented by white coloring. The rightmost panel is a generalized model of the ClpA D2 AAA+ hexamer docking into the ClpP interface. (**B**) Rotary translocation model with clockwise around-the-ring ATP hydrolysis and IGL loop release and rebinding (Ripstein *et al*., 2020; Lopez *et al*., 2020). When subunit S1 is highest in the spiral, ATP hydrolysis releases the IGL loop of subunit S5 and the AAA+ ring rotates clockwise with respect to ClpP. During rotation, subunit S6 moves to the top in of the spiral, and the IGL loop of subunit S5 takes a clockwise “step” and rebinds to the adjacent empty ClpP cleft. Repetition of this sequence of ATP hydrolysis and IGL loop release and rebinding results in rotary motion of the AAA+ ring with respect to the ClpP ring.

In both ClpAP and ClpXP, subunits of the AAA+ enzymes form a shallow spiral, as is observed in most AAA+ proteases and protein-remodeling motors (Puchades *et al*., 2020). In heterohexameric motors in which the positions of unique subunits can be determined, structures with different subunits occupying the highest and lowest positions are observed, suggesting that dynamic rearrangement of the spiral is part of the mechanical cycle used by this enzyme family (de la Peña *et al*., 2018; Dong *et al*., 2019). In one model for this cycle, an enzyme power stroke starts when the second lowest subunit (S5 at the beginning of the cycle) in the spiral hydrolyzes ATP, resulting in a sequential rearrangement that moves this subunit, higher subunits, and bound substrate one position down in the spiral, while the lowest subunit (S6) disengages from the substrate and moves to the top of the spiral (Figure 1B; Puchades *et al*., 2020). Intriguingly, the unoccupied ClpP clefts in ClpAP and ClpXP are always located between the IGL/IGF loops of the second lowest and lowest subunits within the spiral (S5 and S6 in Figure 1A). If subunits in the ClpA or ClpX hexamers pass sequentially through each position in the spiral and the empty clefts in ClpP are always between two specific subunits, then the AAA+ ring should rotate with respect to the ClpP ring during protein translocation (Figure 1B; Ripstein *et al*., 2020; Lopez *et al*., 2020).

Here, we test the effects of preventing rotation by covalently crosslinking multiple IGL loops of ClpA to ClpP. These crosslinked complexes retain substantial proteolytic activity against unfolded and metastable native substrates but display defects in degrading more stably folded proteins. We conclude that rotation of ClpA with respect to ClpP is not required for translocation or unfolding but that some freedom of movement at the ClpA-ClpP interface is likely to be important for optimizing mechanical activities.

## Results and Discussion

### Crosslinking ClpA to ClpP

The penultimate C-terminal residue of ClpP forms part of the binding cleft for the IGL loops of ClpA and appears close enough to allow chemical crosslinking between the IGL loops of ClpA and the C-terminal region of ClpP (Figure 2A) (Lopez *et al*., 2020). We introduced an E613C mutation into the IGL loop of a cysteine-free ClpA^‡^ variant (C47S, C203S, C243S) to generate ^E613C^ClpA^‡^. We also engineered a ClpP variant with the two wild-type cysteines removed by C91V and C113A mutations and inserted a new cysteine after the C-terminal residue (Asn^193^) to generate ClpP^+C^. For crosslinking, we mixed ^E613C^ClpA^‡^ with ClpP^+C^ in the presence of a homobifunctional cysteine crosslinker and then separated the covalently joined ^E613C^ClpA^‡^–ClpP^+C^ complexes away from uncrosslinked species by gel filtration (Figure 2B). Quantification following SDS-PAGE revealed that 90 ± 1% of the ClpA in the gel-filtered pool was present as a crosslinked A– P species and 10 ± 1% was uncrosslinked (Figure 2C Lane 7). This pool also contained uncrosslinked ClpP and crosslinked ClpP dimers. As ∼90% of the ClpA protomers in the pool were crosslinked to ClpP, the vast majority of ClpA hexamers should contain one or more crosslinked A–P subunits (>99.99%, assuming independent crosslinking), and ∼98% of the hexamers in this pool should contain four, five, or six ClpA subunits crosslinked to ClpP.

**Figure 2.**
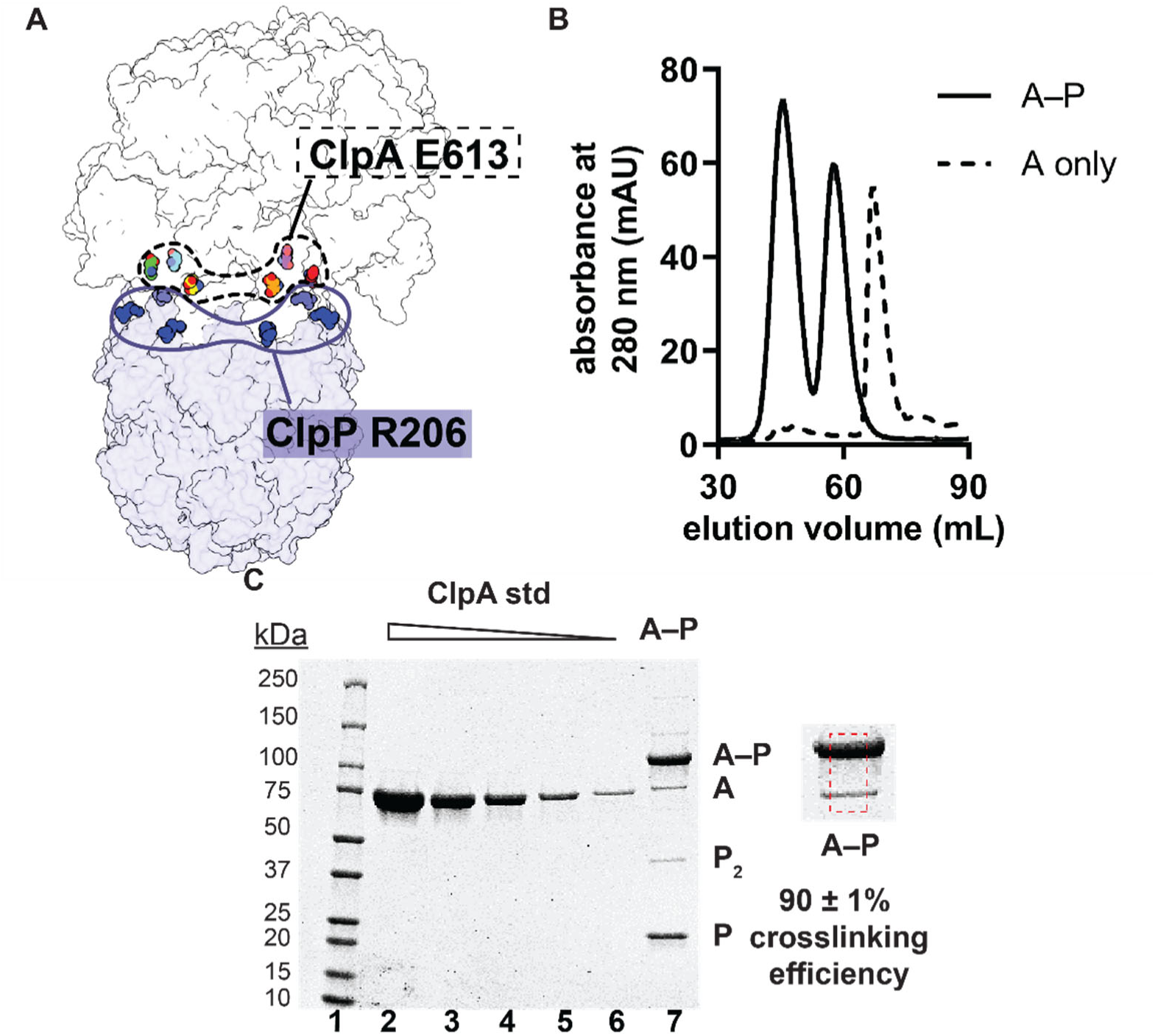
ClpA–ClpP crosslinking and purification (**A**) Proximity of E613 in ClpA to the penultimate residue (R192) of mature ClpP (PDB 6UQO). (**B**) Gel-filtration chromatograms of ClpA only (dashed line) or ClpA–ClpP following crosslinking (solid line). The first peak in the A–P sample (∼45 mL) corresponds to crosslinked A–P. The second peak (∼60 mL) corresponds to uncrosslinked ClpP^+C^. (**C**) Reducing SDS-PAGE gel. Lanes 1-6 are MW standards or different concentrations of purified ^E613C^ClpA^‡^. Lane 7 is a fraction from the A–P pool following gel filtration. The shift in molecular weight from A to A–P denotes a ClpP monomer (∼23 kDa) crosslinked to a ClpA monomer (∼83 kDa). The dashed red box is a zoomed-in view of lane 7 used to calculate crosslinking efficiency of ^E613C^ClpA^‡^ to ClpP^+C^. Crosslinking efficiency was calculated as the mean ± 1 SD of three independent replicates.

### Crosslinked complexes degrade model substrates

To test whether ClpA rotation relative to ClpP is required for ATP-fueled proteolysis, we measured ATP hydrolysis as well as degradation of several model substrates by the A–P pool compared to an A•P control consisting of assembled but uncrosslinked ^E613C^ClpA^‡^ and ClpP^+C^. ATP-hydrolysis by the A–P pool occurred at a rate of 412 ± 40 min^-1^ enz^-1^, whereas this rate was 1034 ± 40 min^-1^ enz^-1^ for the A•P control. Notably, the A–P pool degraded multiple substrates, including the N-terminal domain of the phage λ cI repressor tagged by ssrA (λ cI^N^-ssrA; Gottesman *et al*., 1998), ^cp7^GFP-ssrA (Nager *et al*., 2011), ^5-IAF^titin^I27^-ssrA (Kenniston *et al*., 2003; Iosefson *et al*., 2015), and FITC-casein (Twining, 1984; Thompson *et al*., 1994). Under the conditions of these assays, the A–P pool degraded the folded substrates λ cI^N^-ssrA and ^cp7^GFP-ssrA at rates that were 34 ± 3% and 31 ± 4%, respectively, of the A•P control, and degraded unfolded ^5-IAF^titin^I27^-ssrA and FITC-casein at 44 ± 4% and 97 ± 7%, respectively, of the control rates (Figure 3A-B). The rate of degradation of FITC-casein by the A–P pool was reduced ∼6-fold when ATPγS was substituted for ATP (Figure 3C), indicating that robust degradation of this molten-globule substrate requires ATP hydrolysis. We also determined steady-state kinetic parameters for degradation of ^cp7^GFP-ssrA by the A–P pool and A•P control (Figure 3D). Compared to the control reaction, Vmax was ∼50% and *K*M was ∼3-fold higher for degradation of this substrate by the A–P pool. This reduction in Vmax for the A–P pool was roughly comparable to its reduction in ATP-hydrolysis activity, suggesting that slower degradation of folded substrates by the A–P pool may result from slower ATPase activity. Thus, our results show that multiple crosslinks between ClpA and ClpP in the A–P pool cause modest slowing of the rates of ATP hydrolysis and protein degradation, particularly for native substrates. However, these crosslinks do not prevent the protein unfolding or translocation steps required for effective proteolysis.

**Figure 3.**
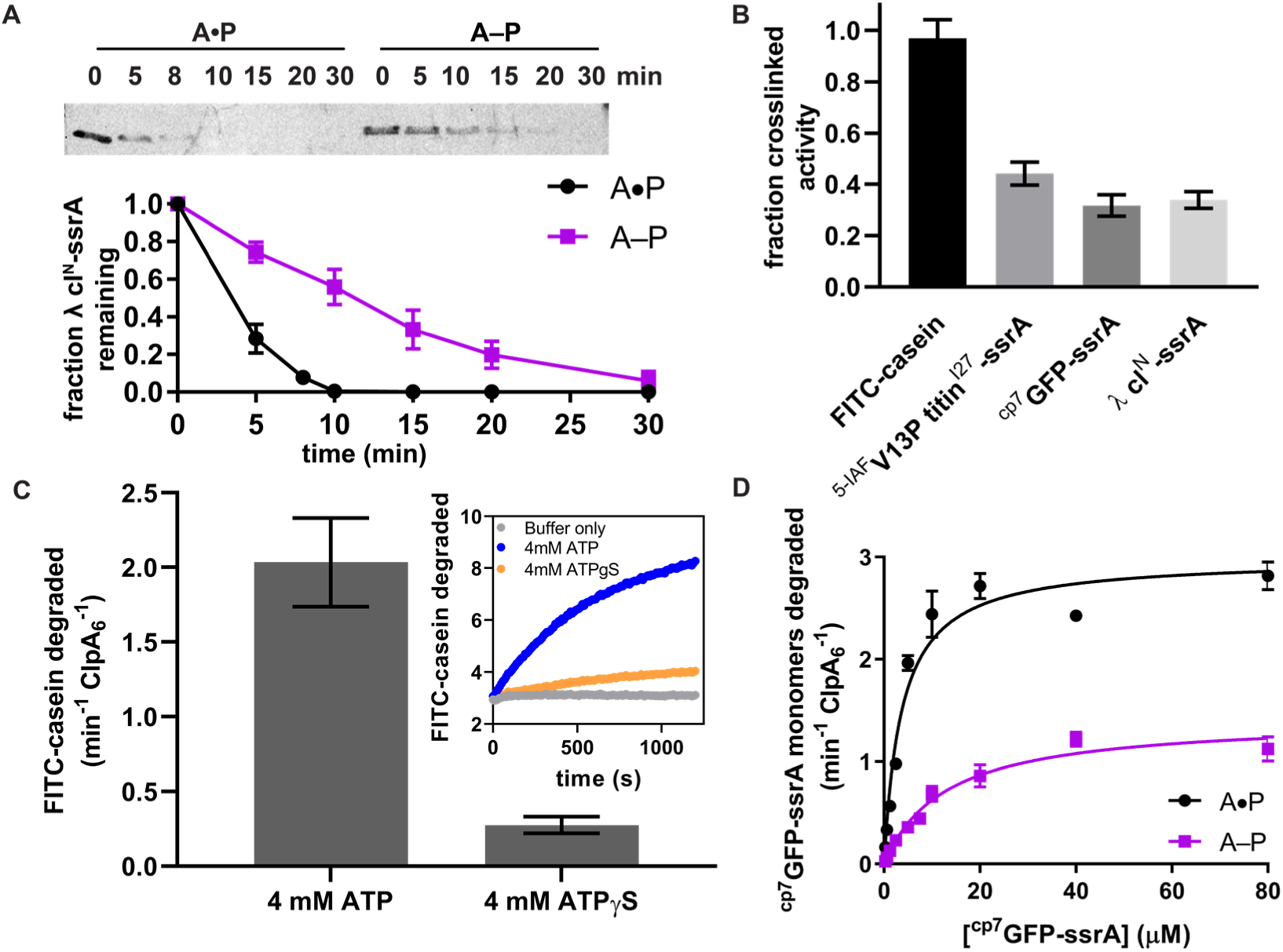
ClpA–ClpP substrate degradation. (**A**) Top, SDS-PAGE assay of the kinetics of λ cI^N^-ssrA degradation by the uncrosslinked A•P control (containing equivalent concentrations of ^E613C^ClpA^‡^ and ClpP^+C^ to the crosslinked pool) and the A–P pool Bottom, quantification of λ cI-ssrA degradation. Values are means ± 1 SD (n ≥ 3). (**B**) Degradation of substrates of varying thermodynamic stability (18µM FITC-casein, 5µM ^5-IAF^titin^V13P^-ssrA, 20µM ^cp7^GFP-ssrA, 15 µM λ cl^N^-ssrA by the purified A–P pool. Fraction crosslinked activity was calculated by normalizing to the activity of the A•P sample. Values are means ± 1 SD (n ≥ 3). (**C**) Degradation of FITC-casein (18 µM) by the purified A–P pool in the presence of ATP or ATPγS. FITC-casein degraded was quantified by normalizing the relative fluorescence units to the total FITC-casein degraded upon porcine elastase addition at the endpoint of the assay and subtracting the contributions of photobleaching from the buffer-only control. Values are means ± 1 SD (n ≥ 3). The inset shows representative kinetics of FITC-casein degradation assay. (**D**) Michaelis-Menten analysis of ^cp7^GFP-ssrA degradation kinetics by the A–P pool and A•P control. Values are means ± 1 SD (n ≥ 3). For the A–P pool, Vmax was 1.4 ± 0.07 min^-1^ ClpA6^-1^, *K*M was 13 ± 1.6 µM, and *R*^2^ was 0.96; for the A•P control, the Vmax was 3.0 ± 0.10 min^-1^ ClpA6^-1^, *K*M was 3.7 ± 0.5 µM, and *R*^2^ was 0.96, where the errors are those of non-linear least-squares fitting to the Michaelis-Menten equation.

### Activity of crosslinked complexes is consistent with a reciprocating mechanism

The sequential hand-over-hand mechanism posits that each subunit in a AAA+ hexamer cycles through each spiral position during processive translocation (Puchades *et al*., 2020). Further, in ClpAP cryo-EM structures, a single empty ClpP cleft is always located between the clefts occupied by the IGL loops of the lowest and second-lowest subunits (S6 and S5 in Figure 1A); when a second empty cleft is observed, it results from disengagement of the IGL loop of the S5 subunit (Lopez *et al*., 2020). Very similar structural results have been observed for ClpXP complexes (Fei *et al*., 2020; Ripstein *et al*., 2020) If ClpA or ClpX subunits cycle through each spiral position, then the empty ClpP cleft would rotate relative to the unfoldase (Figure 4A-B). This model predicts that crosslinking the AAA+ unfoldase to ClpP should abolish protein degradation, as it would stop the rotation and therefore the sequential movement of ClpA/ClpX subunits through each position in the spiral. However, when we prevent ring-ring rotation by crosslinking ClpA to ClpP, substantial rates of degradation of native and denatured protein substrates are still observed. ClpXP complexes in which one IGF loop is crosslinked to ClpP can also degrade folded and unfolded substrates, albeit at lower rates than uncrosslinked controls (Bell, 2020). The high degree of crosslinking in our ClpAP experiments, where 98% of complexes contain at least four covalent crosslinks between ClpA and ClpP, and ∼50% of complexes contain six crosslinks, would be expected to hinder each ClpA subunit from adopting each position in the spiral. Moreover, in approximately half of the crosslinked enzymes, it would not be possible to have two empty ClpP clefts. Hence, the proposal that this intermediate is a requisite step in translocation (Lopez *et al*., 2020) is inconsistent with our results. We conclude that rotation of the ClpA or ClpX rings with respect to the ClpP ring is not required for degradation. Related AAA+ proteases in the FtsH/Yme1/Agf3l2 and Lon families have AAA+ and peptidase modules that are genetically tethered, and therefore also must operate without rotation between the unfoldase and protease components (Glynn, 2017).

**Figure 4.**
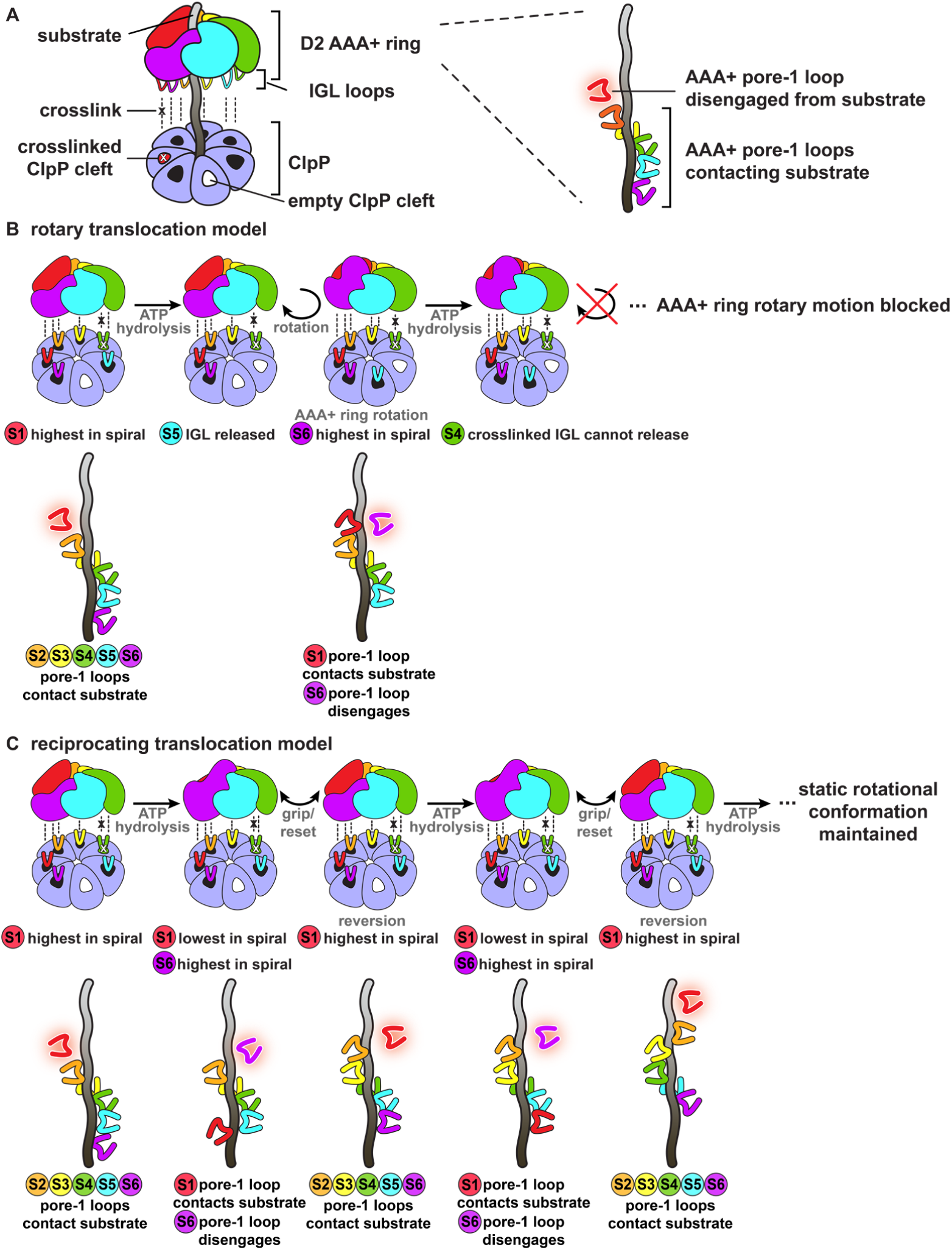
Models for substrate translocation by ClpAP consistent with at least one crosslinked IGL loop. In all panels, ClpA D2 AAA+ subunits are colored as in Figure 1, where subunit S1 is colored red and located at the highest position in the spiral at the beginning of the translocation cycle. (**A**) Generalized model of complexes of ClpA D2 AAA+ hexamer crosslinked to ClpP. The rightmost panel represents the position of pore-1 loops from the corresponding colored AAA+ subunit contacting the substrate during the translocation cycle. The pore-1 loop disengaged from the substrate is indicated by the red border. (**B**) Subunits in the hexamer move sequentially through each position in the spiral, as changes in IGL loop conformations result in movement of the empty ClpP cleft. (**C**) Reciprocating translocation model in which a subset of IGL loops maintain static interactions with ClpP. Only the highest and lowest subunits in the spiral change positions.

A reciprocating mechanism in which the AAA+ unfoldase maintains a static rotational conformation relative to ClpP during substrate processing could, in principle, account for our results as well as ClpAP and ClpXP structures. For example, as shown in Figure 4B, the highest subunit in the spiral could move to the lowest position, dragging the substrate through a ∼6-residue translocation step, while the lowest subunit moves to the highest position (see movie S4 in Fei *et al*., 2020). Other subunits could then grip the substrate as the subunit in the lowest position in the spiral reverts to its original position for the next power stroke (Figure 4C, lower). This model requires just two subunits to dynamically exchange positions along the seam interface of the hexamer. Crosslinking could hinder but not prevent such motion, explaining the reduced activity of our A–P variants. This general model is also consistent with experimental measurements of the basic translocation step size for ClpXP and ClpAP (Aubin-Tam *et al*., 2011; Maillard *et al*., 2011; Miller *et al*., 2013; Sen *et al*., 2013; Cordova *et al*, 2014; Olivares *et al*., 2014; 2017). ClpA and ClpX variants with mixtures of ATPase active and inactive modules support significant rates of ClpAP/ClpXP degradation of protein substrates (Martin et al., 2005; K. L. Zuromski and T.A. Baker, personal communication, May 2020). Hence, ATP hydrolysis in any one of multiple subunits in the AAA+ spiral may suffice to drive a power stroke.

## Materials and Methods

### Proteins

The gene encoding *E. coli* ClpP^+C^ was generated using PCR mutagenesis, and the corresponding protein was purified by established protocols and stored in buffer containing 0.5 mM dithiothreitol (DTT) (Martin *et al*., 2005). Wild-type ClpP was purified by established protocols (Kim *et al*., 2001). The plasmid for ^E613C^ClpA^‡^ was generated by PCR mutagenesis of *E. coli* ClpA^ΔC9^ fused to the 3’-end of His6SumoFLAG cloned into pET23b (Novagen). ClpA was overexpressed in T7Express (NEB), purified by Ni-NTA chromatography, followed by ULP-1 cleavage and SP-Sepharose cation exchange, and stored in 50 mM HEPES-KOH, pH 7.5, 300 mM NaCl, 20 mM MgCl2, 10% glycerol, 2 mM TCEP. The ^cp7^GFP-ssrA and V13P titin^I27^-ssrA proteins were purified according to established protocols (Nager *et al*., 2011, Kenniston *et al*., 2003). V13P titin^I27^-ssrA was labeled with 5-iodoacetamidofluorescein (5-IAF) for fluorescent assays by an established protocol (Iosefson *et al*., 2015). The plasmid for λ cI^N^-ssrA was generated by PCR mutagenesis of bacteriophage λ cI (amino acids 1-93) fused to the 3’-end of His6Sumo cloned into pET23b and appended with the ssrA degron. λ cI^N^-ssrA was purified by Ni-NTA chromatography, followed by ULP-1 cleavage, Ni-NTA to remove the H6SUMO fragment, Mono-Q anion exchange, and Superdex-75 gel filtration chromatography, and stored in 25 mM HEPES-KOH pH 7.5, 150 mM NaCl, 10% glycerol, 1 mM DTT.

### Crosslinking ClpA to ClpP

^E613C^ClpA^‡^ (4 μM) and ClpP^+C^ (9.6 μM) were mixed in a total volume of 2.5 mL and desalted into 50 mM HEPES-KOH, pH 7, 300 mM NaCl, 20 mM MgCl_2_, 10% glycerol, 5 mM EDTA over a Sephadex G-25 PD-10 column (GE Healthcare). After diluting to a final volume of 5 mL, crosslinking was initiated by addition of 5 mM ATPγS and 200 μM bismaleimidoethane (BMOE; Thermo Fisher) and allowed to proceed at room temperature for 45 min. The reaction was quenched by addition of 50 mM DTT at room temperature for 20 minutes before purification by Superdex-200 gel filtration chromatography in ClpA storage buffer. Crosslinked species were measured by quantifying Coomassie-stained SDS-PAGE gel bands relative to ^E613C^ClpA^‡^ standards. The area under the curve (AUC) corresponding to density of the crosslinked and uncrosslinked species were quantified by ImageQuant (GE Healthcare) after scanning Coomassie-stained SDS-PAGE gels using a Typhoon FLA 9500 (GE Healthcare). Crosslinking efficiency was measured in three independent replicates, and calculated as:

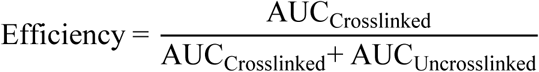

### Biochemical Assays

ATP-hydrolysis assays were performed using an NADH-coupled assay, as described previously (Martin *et al*., 2005), at 30 °C in Buffer HO (25 mM HEPES-KOH, pH 7.5, 300 mM NaCl, 20 mM MgCl2, 10% glycerol, 2 mM TCEP) with 5 mM ATP and 0.25 µM ^E613C^ClpA^‡^ hexamer and 0.75 µM ClpP14 for the A•P control or 0.25 µM ^E613C^ClpA^‡^– ClpP^+C^ hexamer for the A–P pool. All ClpAP degradation reactions were performed at 30 °C in Buffer HO with an ATP regeneration system of 4 mM ATP, 50 μg/mL creatine kinase (Millipore-Sigma), and 5 mM creatine phosphate (Millipore-Sigma). Degradation of ^cp7^GFP-ssrA (20 µM monomer for the “fraction crosslinked activity”) containing 0.25 µM ^E613C^ClpA^‡^ hexamer and 0.75 µM ClpP14 for the A•P control or 0.25 µM ^E613C^ClpA^‡^– ClpP^+C^ hexamer for the A–P pool was monitored by loss of substrate fluorescence (excitation 467 nm; emission 511 nm), measured using a SpectraMax M5 plate reader (Molecular Devices) (Nager *et al*., 2011). Degradation of FITC-casein (18 µM, Sigma-Aldrich) containing 0.25 µM ^E613C^ClpA^‡^ hexamer and 0.75 µM ClpP14 for the A•P control or 0.25 µM ^E613C^ClpA^‡^–ClpP^+C^ hexamer for the A–P pool was monitored by increase in fluorescence (excitation 340 nm; emission 520 nm); to determine the endpoint of complete FITC-casein degradation, 0.5 µL of 5 mg/mL porcine elastase (Sigma-Aldrich) was added to each well and incubated for 30 min. ClpAP degradation reactions with FITC-casein (18 µM) were performed at 30 °C in Buffer HO with 4 mM ATP or ATPγS (Millipore Sigma). Degradation of ^5-IAF^V13P titin^I27^-ssrA (5 µM) containing 0.2 µM ^E613C^ClpA^‡^ hexamer and 0.75 µM ClpP14 for the A•P control or 0.2 µM ^E613C^ClpA^‡^– ClpP^+C^ hexamer for the A–P pool was monitored by increase in fluorescence (excitation 494 nm; emission 518 nm). Gel degradation of λ cI^N^-ssrA (15 µM monomer) containing 0.2 µM ^E613C^ClpA^‡^ hexamer and 0.4 µM ClpP14 for the A•P control or 0.2 µM^E613C^ClpA^‡^–ClpP^+C^ hexamer for the A–P pool was performed in triplicate by taking samples of each reaction at specific time points, stopped by addition of SDS-PAGE loading dye and boiling at 100 °C before loading on Tris-Glycine-SDS gels. Bands were visualized by staining with colloidal Coomassie G-250 and quantified by ImageQuant (GE Healthcare) after scanning by Typhoon FLA 9500 (GE Healthcare). The fraction of λ cI^N^-ssrA remaining was calculated by dividing the intensity of this band at a given time point by the density at time zero, after normalization by the creatine kinase density. All experiments were performed in at least three independent replicates and values reported were calculated as the mean ± 1 SD of independent replicates.

## Acknowledgments

We are grateful to X. Fei (MIT) for helpful advice on preparation of crosslinked samples. This work was supported by NIH grants GM-101988 (RTS), AI-016892 (RTS, TAB), and the Howard Hughes Medical Institute (TAB). TA Bell and SK were supported by NIH training grant 5T32GM-007287; SK and KLZ were supported by NSF grant GRFP 1122374.

